# Enhanced flavour profiles through radicicol induced genomic variation in *S. pastorianus* lager yeast

**DOI:** 10.1101/2022.05.17.491830

**Authors:** Roberto de la Cerda Garcia-Caro, Georgia Thompson, Penghan Zhang, Karsten Hokamp, Fiona Roche, Silvia Carlin, Urska Vrhovsek, Ursula Bond

**Affiliations:** Department of Microbiology, Moyne Institute, School of Genetics and Microbiology, Trinity College Dublin, Dublin 2, Ireland; Metabolomic Unit, Food Quality and Nutrition Department, Research and Innovation Centre, Edmund Mach Foundation, Via E.Mach 1, 38010 S.Michele all’Adige, Italy; Department of Genetics, Smurfit Institute, School of Genetics and Microbiology, Trinity College Dublin, Dublin 2, Ireland

## Abstract

The yeasts, *Saccharomyces pastorianus*, are hybrids of *Saccharomyces cerevisiae* and *Saccharomyces eubayanus* and have acquired traits from the combined parental genomes such as ability to ferment a range of sugars at low temperatures and to produce aromatic flavour compounds, allowing for the production of lager beers with crisp, clean flavours. The polyploid strains are sterile and have reached an evolutionary bottleneck for genetic variation. Here we describe an accelerated evolution approach to obtain lager yeasts with enhanced flavour profiles. As the relative expression of orthologous alleles is a significant contributor to the transcriptome during fermentation, we aimed to induce genetic variation by altering the *S. cerevisiae* to *S. eubayanus* chromosome ratio. Aneuploidy was induced through the temporary inhibition of the cell’s stress response and strains with increased production of aromatic amino acids via the Shikimate pathway were selected by resistance to amino acid analogues. Genomic changes such as gross chromosomal rearrangements, chromosome loss and chromosome gain were detected in the characterised mutants, as were Single Nucleotide Polymorphisms in *ARO4*, encoding for DAHP synthase, the catalytic enzyme in the first step of the Shikimate pathway. Transcriptome analysis confirmed the upregulation of genes encoding enzymes in the Ehrlich pathway and the concomitant increase in the production of higher alcohols and esters such as 2-phenylethanol, 2-phenylethyl acetate, tryptophol, and tyrosol. We propose that the plasticity of polyploid *S. pastorianus* genomes is an advantageous trait supporting opportunities for genetic diversity in otherwise sterile strains.

**Significance Statement:** Lager beer is the product of fermentations conducted with *Saccharomyces pastorianus*, which are hybrids of *Saccharomyces cerevisiae* and *Saccharomyces eubayanus*. A quintessential property of lager beers is the distinctive flavours produced during fermentation. Hybrids are sterile and have reached an evolutionary bottleneck. Finding ways to introduce genetic variation as a means of enhancing the flavour profiles is a challenge. Here, we describe an approach to introduce genetic variation by inducing aneuploidy through the temporary inhibition of the cell’s stress response. Strains with an enhanced flavour production were selected by resistance to amino acid analogues. We identified genomic changes and transcriptome analysis confirmed the upregulation of genes in the Ehrlich pathway which is responsible for the production of flavour compounds.

## Introduction

The lager yeasts, *Saccharomyces pastorianus*, are natural hybrids of *Saccharomyces cerevisiae* and *Saccharomyces eubayanus* that emerged during the Middle Ages in Central Europe (1). The hybridisation events converged the high fermentative rates of *S. cerevisiae* with the cryotolerance properties of *S. eubayanus*. The evolution of *S. pastorianus* has been influenced by human laws and practices (2). In modern day breweries, fermentations with *S. pastorianus* are conducted at temperatures between 7-13°C, creating a beer with crisp, clean flavours that is the preferred beverage of most customers (3).

The complex polyploid genomes of *S. pastorianus* strains underpin the physiological outcome of the fermentation. The strains are classified into two broad groups, Group I and Group II that differ in the relative proportion of *S. cerevisiae* and *S. eubayanus* DNA content. Group I, or *Saaz* strains, are typically triploid in DNA content, retaining all the parental *S. eubayanus* chromosomes but have lost many *S. cerevisiae* chromosomes. The Group II, or *Frohberg* strains, are mainly tetraploid in DNA content, containing approximately 2n *S. cerevisiae* and 2n *S. eubayanus* genome content (4-6). Both groups display chromosomal aneuploidy with chromosome numbers ranging from one to six (5, 7). In addition to the parental chromosomes, *S. pastorianus* strains contain several hybrid chromosomes containing both *S. cerevisiae* and *S. eubayanus* genes that resulted from recombination, at precise locations, between the parental chromosomes. Some of the recombination breakpoints are located within coding regions, creating a set of hybrid genes unique to lager yeasts (1, 6, 8, 9).

The presence of orthologous alleles, emanating from different parental chromosomes, creates a complex network of gene transcription. Superimposed on the presence of orthologous alleles are complexities of gene dosage due to the aneuploid nature of the genomes. The resultant steady state pool of mRNAs has the potential to create a complex proteome with the potential to affect the stoichiometry of *S. cerevisiae* and *S. eubayanus* proteins within protein complexes (10) and to influence the final cellular physiology and the fermentation properties of *S. pastorianus* strains (11).

We recently analysed the patterns of gene expression under fermentation conditions in representative Group I and Group II lager yeasts and showed that the steady state levels of *S. cerevisiae* and *S. eubayanus* orthologous alleles are directly correlated with the copy number of the genes (12). Furthermore, we observed that *S. cerevis*iae and *S. eubayanus* alleles differentially contribute to metabolic processes in the cell, and in some cases, contribute exclusively to specific gene ontologies. Flavour and aroma profiles impart unique characteristics to lager beers and arise from the production of higher alcohols and acetate esters as secondary metabolites from the catabolism of aromatic, branched-chain, and sulphur-containing amino acids via the Ehrlich pathway (3, 13). Additionally, fatty-acid esters produced from the esterification of Acyl-CoA with ethanol add to the final organoleptic profile of the fermentation.

*S. pastorianus* hybrids are generally sterile, rarely producing spores and what spores are produced have low viability, therefore the existing hybrid strains have limited genetic variation. Hybrid sterility is most likely a consequence of a lack of normal mitotic recombination or meiotic cross-over events (14). Increasing genetic diversity is challenging, but several approaches such as laboratory evolution, traditional mutagenesis and interspecific hybridisation have been used to generate new strains with phenotypes relevant to brewing (15-21). Genetic variation within hybrids can also be manifested through chromosomal recombination and changes in aneuploidy (22-24).

Here we have used an accelerated evolution approach to generate novel strains of *S. pastorianus* strains with varied flavour profiles. As the relative expression of orthologous alleles was shown to be a significant contributor to the final gene expression pattern during fermentation, we aimed to alter the *S. cerevisiae* to *S. eubayanus* chromosome ratio in lager yeast strains. We induced chromosomal aneuploidy and gross rearrangements by exposing cells to Heat Shock Thermal Stress (HSTS) or to the macrocyclic antibiotic radicicol, an inhibitor of Hsp90p (22, 23). By coupling the stress treatment to the ability of mutants to grow in the presence of amino acid analogues of phenylalanine, we selected for variants with increased production of higher alcohols and esters produced via the Ehrlich pathway.

## Results

### Mutant selection by heat stress and radicicol treatment

Two representative *S. pastorianus* strains, the Group I strain CBS1538, and the Group II strain W34/70 were chosen for analysis. The strains display similar organoleptic volatile profiles at the end of fermentation, yet each displays unique flavour profiles, with CBS1538 producing higher levels of ethyl butyrate and ethyl hexanoate that impart tropical fruit flavours and W34/70 producing higher levels of methionol, which produces a “worty” note in fresh beers (*SI Appendix*, Fig. S1). Due to the differences in copy numbers of *S. cerevisiae* (*Sc*) and *S. eubayanus* (*Se*) chromosomes, each strain possesses a unique gene expression profile with *Sc* and *Se* alleles differentially contributing to the steady state pool of mRNAs encoding proteins associated with different metabolic processes (12). For certain metabolic activities, active during the fermentation process, *Sc* or *Se* alleles were observed to exclusively encode for proteins within specific biochemical pathways.

With the aim of introducing genetic variation, the strains were exposed to varying temperatures or concentrations of radicicol (*SI Appendix*, Table S1) and following the treatment were plated onto minimal media agar plates containing amino acid analogues of phenylalanine, either β-2-thienylalanine (B2TA) or ρ-fluorophenylalanine (PFPA). Such amino acid analogues select strains with impaired negative feedback inhibition on the committed step in the Shikimate pathway, increasing the biosynthesis of aromatic amino acids and the flux through the Ehrlich pathway towards the production of higher alcohols and esters. A total of 96 mutants were isolated.

Heat shock treatment followed by selection on B2TA produced only clones from strain W34/70. Likewise, no clones were produced by radicicol treatment followed by B2TA selection in either strain (*SI Appendix*, Table S1). The isolation rate following heat shock treatment for W34/70 and CBS1538 were 1.86 × 10^-7^ and 1.08 × 10^-7^ respectively. Isolation rates for radicicol treatment were 7.66 × 10^-7^ for W34/70 and 7.33 × 10^-7^ for CBS 1538. No clones were obtained if the parental strains were plated on minimal medium with amino acid analogues without any treatment.

### Characterisation of mutants

Twenty-two random mutants were selected for growth characterization and aromatic profile analysis in a small-scale fermentation. The mutants were grown in minimum medium containing different concentrations of the amino acid analogues (Fig. 1A) and under different growth conditions (Fig. 1B). The mutant strains grew at rates similar to the parental strains in minimal medium (0.95 ± 0.08) and in rich medium at 20°C (0.96 ± 0.09) and 30°C (0.94 ± 0.11) with some strains, CBS1538 1.1, 1.2, and 11.1, displaying significantly more growth in 30°C in YPDM and strains W34/70 8.4, 9.1 displaying increased growth at 20°C (Fig. 1B). In minimal medium containing phenylalanine as the sole nitrogen source, almost all the strains exhibit a similar growth rate to their respective parental strain (0.8 ± 0.16), except two Group II mutants, WS 7.3 and WS 8.1 that displayed lower growth.

**Figure 1.**
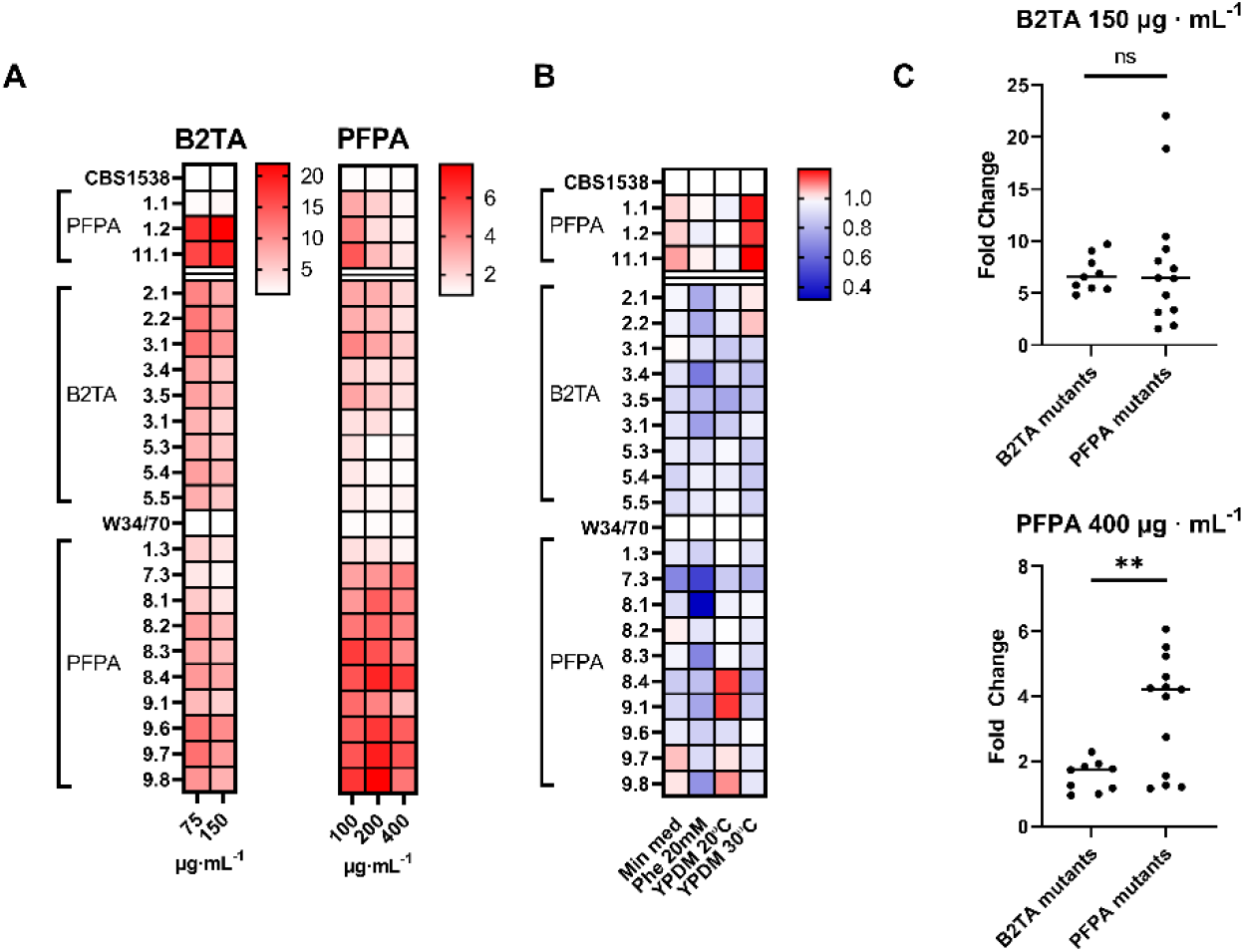
Characterization of the mutant and parental strains in different concentration of amino acid analogues (A) and under different physiological conditions (B). Values correspond to the Area under the Curve (AUC) during 68h of growth and normalized to the respective parental strain which was set at a nominal value of 1.0 (white box). The growth of mutants relative to the wildtype strains are shown in the heat map legends. C. Range and median AUC of mutants selected in 150 mg · mL^-1^ B2TA (top) and 400 mg · mL^-1^ PFPA (bottom). ns, no significant difference; **, p value <0.05

Strains selected for resistance to PFPA displayed cross resistance to B2TA (Fig. 1C). Likewise, B2TA mutants displayed resistance to PFPA, however the growth rate of such mutants was significantly lower than the PFPA mutants (Fig. 1C).

To compare the aroma profiles of the mutants, small-scale fermentations were carried out and the volatile profiles were examined by semi-quantitative GC-MS analysis at the end of the fermentation (Fig 2), with a specific aim to identify mutants with increased production of higher alcohols and esters produced by the amino acid phenylalanine. Overall, the aromatic profiles produced by the mutants are similar to the parental strains, however some mutants showed distinctive changes. Eight strains (CBS15381.2, 11.1, W34/70 1.3, 2.2, 3.1, 5.5, 9.1 and 9.7 produced higher levels of 2-phenylethanol and seven strains (CBS1538 11.1, W34/70 1.3, 2.2, 3.1, 8.2, 9.1 and 9.7) produced higher levels of 2-phenylethyl acetate compared to the parental strains. Based on the combined data for resistance to amino acid analogues, growth characteristics and flavour profiles, two strains, namely CBS1538 11.1 and W34/70 9.7, both obtained from radicicol treatment and selected on PFPA, were selected for further analysis.

**Figure 2.**
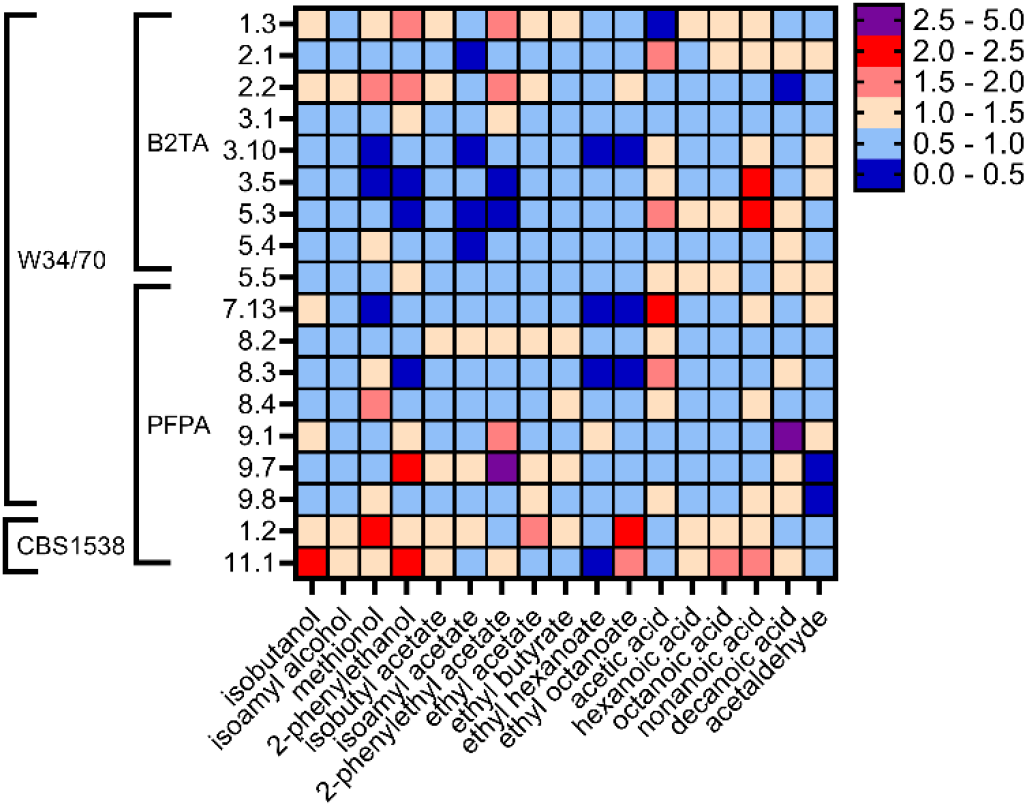
Characterization of the flavour profile of the mutants in small-scale fermentations. Values are normalized by the values of the respective parental strain which was set at 1.0. Colour legend is to the right of heat map.

### Confirmation of the phenotypes of the mutant strains

Growth in liquid minimal medium in the presence of the amino acid analogues confirmed the resistance of the mutants to the amino acid analogues. Mutant 9.7 was resistant up to a concentration of 400 µg · mL^-1^ of PFPA while mutant 11.1 is more sensitive displaying a maximum resistance at 100 µg · mL^-1^ (Fig. 3A).

**Figure 3.**
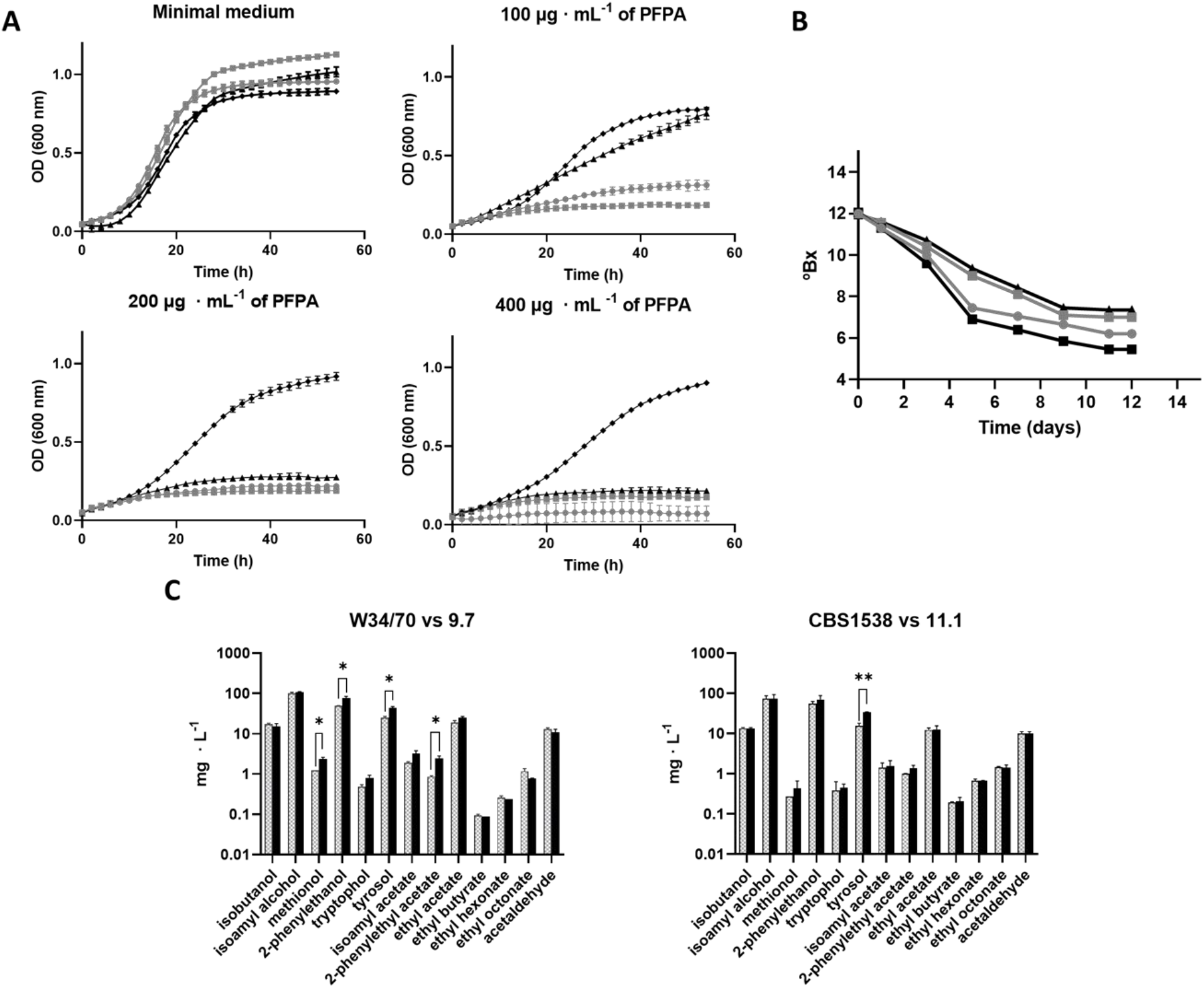
Selected mutants are resistant to the amino acid analogue PFPA and show different fermentative qualities. A) Growth characterization of mutant and WT strains in different concentrations of PFPA. B) Fermentation profiles of the selected strains in 12°Bx wort at 13°C in 3L cylindrical tubes. For A and B, CBS1538: grey line with grey squares, 11.1: black line with triangles, W34/70: grey line with circles, and 9.7: black line with diamonds. C) Aromatic profiles of the selected strains. Parental strains, W34/70 and CBS1538, Hatched columns, mutants 9.7 and 11.1, black columns. Error bars represent the standard deviations from the mean of duplicate fermentations. Compounds showing statistical differences between the mutants and the parental strains are indicated by connecting lines, p≤0.01 **; p≤0.05, *.

To quantify and compare the volatile compounds produced in the mutants and the WT strains, large-scale fermentations were carried out (Fig. 3B). The Group II strain W34/70 fermented faster and reached a greater attenuation than the CBS1538 strain. Surprisingly, the mutant 9.7 fermented faster and consumed more sugars compared to its WT strain (Fig. 3B) while the mutant 11.1 fermented slightly slower than its WT strain.

Quantitative GC/MS analysis of the volatile profiles of the mutant and WT strains confirmed the increased production of 2-phenylethanol and 2-phenylethyl acetate in the mutants 9.7 and 11.1 (Fig. 3C). The increase in these compounds in mutant 11.1 was not deemed statistically different than the parent in this experiment with just duplicate samples, however this trend in higher 2-phenylethanol and 2-phenylethyl acetate was observed in several independent experiments. Mutant 11.1 displayed significantly increased levels of tyrosol while tryptophol levels were significantly increased in mutant 9.7. Both higher alcohols are derived from the aromatic amino acids tyrosine and tryptophan, respectively, via the Ehrlich pathway. Mutant 9.7 also showed increased levels of methionol (Fig. 3C).

### Chromosomal changes in the mutant strains

To identify any genetic changes in the mutant strains, both the mutant and parental strains were sequenced *de novo* and mapped to the annotated and fully assembled reference genome *S. pastorianus* 1483 (Group II strain) as well as to a newly generated *S. pastorianus* “combined” genome, assembled from the parental reference genomes *S. cerevisiae* and *S. eubayanus*. Both approaches yielded comparable results, however since the *S. pastorianus* 1483 genome lacked some information for *S. cerevisiae* genes on chromosomes III and VII, the data from mapping to the combined parental genomes was used to identify any changes in copy number in mutant strains (Fig. 4A).

**Figure 4.**
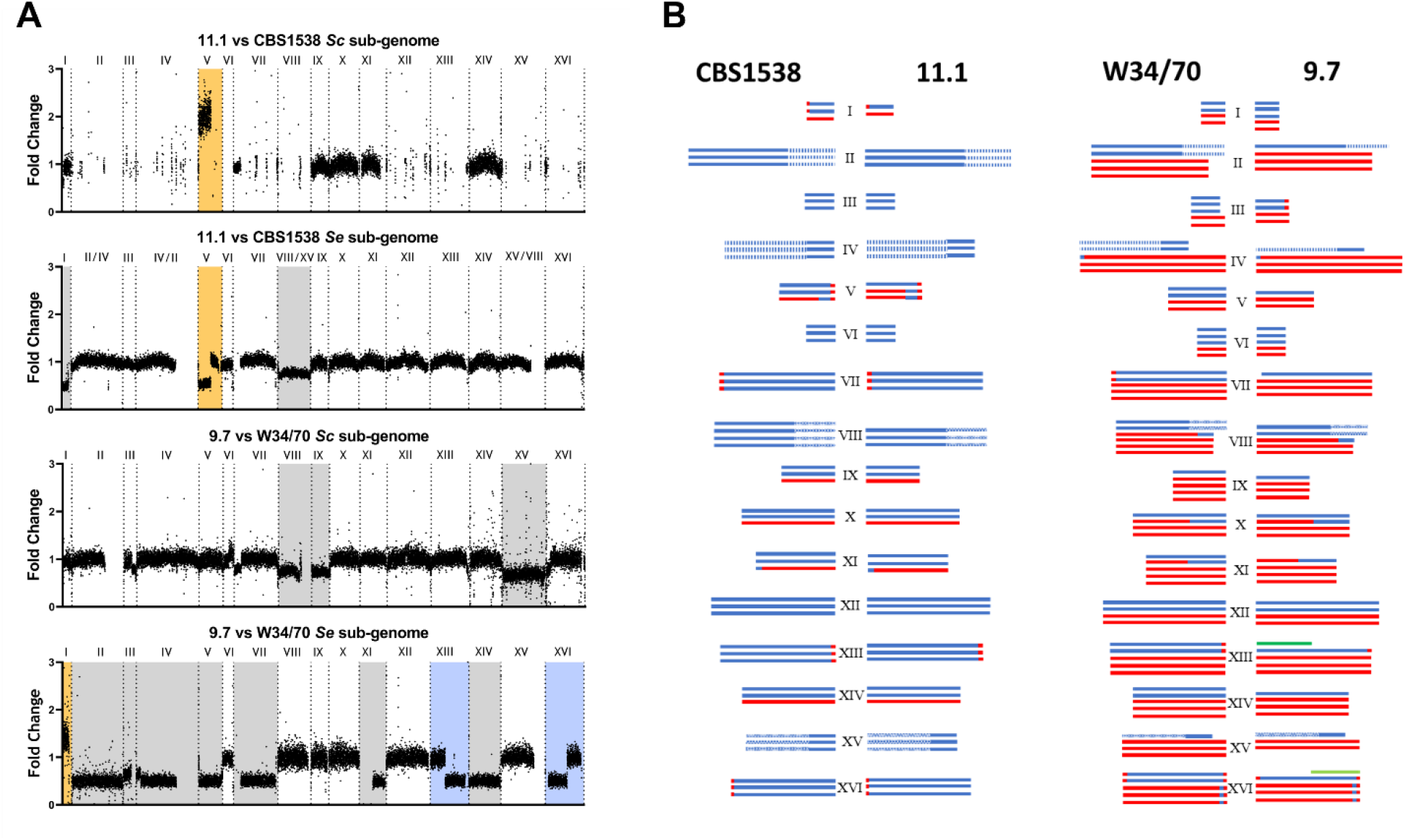
Chromosome maps of the parental and mutant strains. A. Read coverage fold change between mutant and WT strains by chromosome and sub-genome. Dotted lines represent the start of each chromosome. Chromosome copy number gain (gold), chromosome loss (grey), and chromosomal translocations (blue). B. Estimated copy number of *S. cerevisiae* (red), *S. eubayanus* (blue), hybrid (red/blue), and translocated (green) chromosomes in CBS1538 and W34/70 and their respective mutant strains; mutant 11.1 and mutant 9.7.

Mutant 9.7 experienced a greater degree of radicicol induced aneuploidy compared to mutant 11.1 (Fig. 4A). We observed chromosome loss or gain in both mutants: mutant 11.1 displayed a loss of *Se* chromosomes I and VIII/XV (Fig. 4B), while mutant 9.7 has reduced copy number of chromosomes *Se* II/IV, III, IV/II, V, VII, XI, and XIV as well as *Sc* chromosomes VIII, IX and XV. There was also a gain in copy number in *Se* I in mutant 9.7 and a gain of one type of hybrid chromosome *Se* V with a concomitant loss of the second type of hybrid Se V in mutant 11.1 (Fig. 4B). Finally, we observed the formation of a new chromosomal translocation between chromosomes *Se* XIII and *Se* XVI (Fig. 4B). Alignment of the translocation regions shows that both chromosomes contain a region with high similarity in an intergenic region containing a Long Terminal Repeat (LTR) similar to Tsu4. The newly created chromosome *Se* XIII/*Se* XVI contains approximately 367kb of *Se*-like chromosome XIII and 350 kb of *Se*-like chromosome XVI. In total, there is a net loss of 9 *Se* and 3 *Sc* chromosomes, a net gain of 1 *Se* chromosome as well as a new translocation in mutant 9.7, while in mutant 11.1 there is a net loss of 2 *Se* chromosomes as well as a copy number change in Hybrid *Se* V. The copy number changes in selected chromosomes in mutant 9.7 were confirmed by qPCR of genomic DNA (*SI Appendix*, Fig. S3).

### Analysis of allelic variants in W34/70

We also analysed the genomes to identify any Single Nucleotide Polymorphism (SNPs) in the mutant strains using the annotated genome of the Group II strain CBS1483 as the reference strain. The SNP analysis confirmed the presence of two different *S. cerevisiae* sub-genomes in W34/70 strain, referred here as *Sc1* and *Sc2* (4). Analysis of allelic frequencies between mutant 9.7 and W34/70 uncovered changes in chromosome copy number in *Sc1* and *Sc2* that were not detected by the chromosome copy number analysis. Specifically, we detected differences in allelic frequencies between mutant 9.7 and W34/70 on *Sc* chromosomes II, IV and XV. Two types of changes in allelic frequencies were observed, namely, changes in the ratio of allelic variants and loss of heterozygosity (LOH) (*SI Appendix*, Fig. S3) We also analysed the genomes to identify any Single Nucleotide Polymorphism (SNPs) that introduced non-synonymous or stop gain codon changes into protein encoding genes in the mutant strains.

Surprisingly, only a small number of SNPs were unique to the mutant strains (*SI Appendix*, Table S2). Just 25 SNPs, which passed all filters and were deemed unambiguous based on chromosome copy numbers were detected in mutant 9.7 (*SI Appendix*, Table S2). Of interest here is Sc*ARO4*, where two of the three copies of the gene have a non-synonymous SNP (S195F). Mutant 11.1 had just two non-synonymous SNPs, including a SNP in Se*ARO4* (D22Y).

### Gene expression changes in the mutant strains

We examined the gene expression patterns in the mutants 9.7 and 11.1 and the parental strains in three experimental conditions, namely, growth in minimal medium and fermentation in wort on Day 2 and Day 4 (*SI Appendix*, S3 Table). Mutant 9.7 has lost its only copy of *Se* chromosome XI but contains a hybrid copy of chromosome XI (Fig. 4B), leaving a total of 166 *Se* genes missing from the mutant. This gene set was excluded from the analysis. The majority of differentially expressed genes in the mutant 9.7 are condition-specific with growth in minimal medium displaying the greatest number of genes (Fig. 5A). Just 15 genes are upregulated, and 87 genes downregulated in the mutant in all three conditions. The upregulated gene set includes the genes *Sc ARO9* and *Sc ARO10* (*SI Appendix*, Table S4 and S5) encoding for aromatic aminotransferase II and phenylpyruvate decarboxylase, respectively, the enzymes responsible for the first two steps of the Ehrlich pathway. Also of note is the upregulation of *MET32*, the global transcriptional regulator of methionine biosynthetic genes.

**Figure 5.**
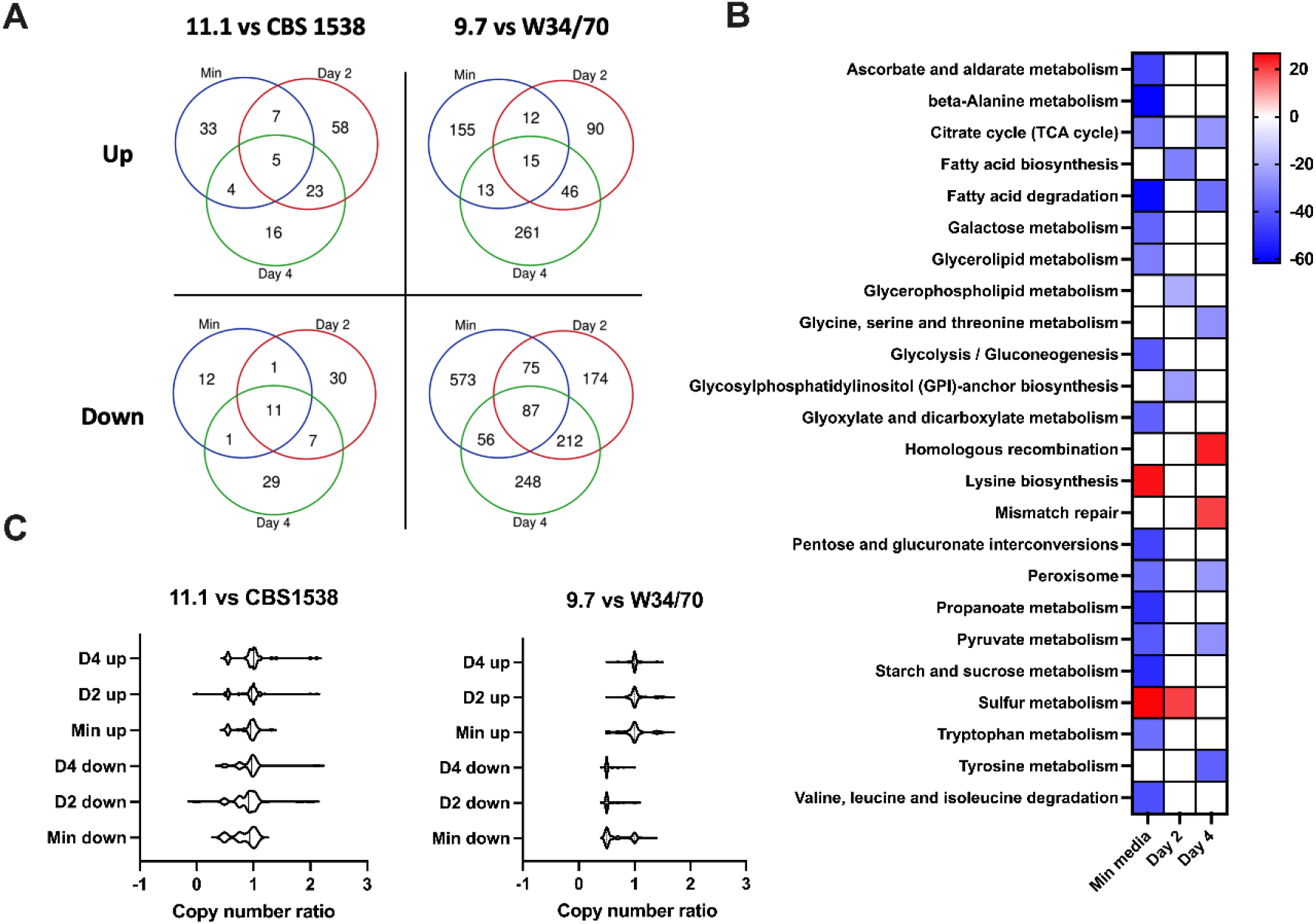
A. Venn diagrams showing numbers significantly (p<0.05) up and downregulated genes common to the three experimental conditions: minimal media, wort Day 2, and wort Day 4. B. Heatmap representing the percentage of genes associated with the listed gene ontologies which are differentially expressed between mutant and wildtype under the different experimental conditions. Upregulated GOs (red), downregulated (blue). White boxes: no significant number of differentially expressed genes under that condition. C. Distribution of copy number ratios (mutant:wildtype) in the different experimental conditions.

Genes involved in sulphur metabolism are upregulated in mutant 9.7 in minimal medium and on Day 2 (*SI Appendix*, Tables S4 and 7) along with the associated sulphur metabolism gene ontology (Fig. 5B). Gene ontologies associated with homologous recombination and mismatch repair were also enriched in mutant 9.7 (Fig. 5B). Downregulated gene ontologies in WS 9.7 in minimal medium include sugar metabolism and amino acid metabolism including beta-alanine metabolism, tryptophan metabolism and valine, leucine, and isoleucine degradation. Day 4 also had downregulated ontologies associated with amino acid metabolism, affecting tyrosine, glycine, serine, and threonine metabolism as well as genes associated with membrane biosynthesis (Fig.5B).

Consistent with the reduced genomic changes in mutant 11.1, we observed fewer differentially expressed genes between the mutant and the parental strain CBS1538. The majority of differentially expressed genes were condition-specific with just 5 genes upregulated and 11 genes downregulated in all three conditions (Fig. 5A). Interestingly amongst the genes upregulated in all three conditions are genes encoding *HSP82, TEF2*, and *MAL32* (*SI Appendix*, Table S6). The heat shock genes *HSP104, HSP30, HSP26* and *SSA4* are also upregulated in mutant 11.1 on Day 2 in wort and genes encoding hexose transporters are upregulated on Days 2 and 4 in wort. The genes *BAP2* and *BAT2* encoding for a branched chain amino acid permease and branched chain amino acid transaminase respectively are upregulated in minimal medium and the ammonium permease *MEP2* is upregulated on Day 2 in wort (*SI Appendix*, Table S6). Amongst the commonly downregulated gene pool are several genes encoding ribosomal proteins. There were no gene ontologies enriched in the up-or down-regulated gene pools in mutant 11.1.

To determine if the gene expression changes in the mutants were due to the observed chromosomal copy number differences in the mutants, the ratio of gene copy number in the mutant and WT strains for up- and down-regulated genes was examined (Fig. 5C). For the 9.7 mutant, there is no change in the gene copy number between the mutant and the parental strain (ratio of copy number = 1.0) for most upregulated genes (Fig. 5C). A small group of genes located in *Se* Chr I have a higher expression rate due to an increase in copy number of this chromosome. However, for the downregulated gene pool, most genes displayed a reduction in the gene copy number (ratio of copy number = 0.5), except for a sub-set of genes in the minimal medium condition. For the mutant 11.1, here again, there is no change in the gene copy number ratio for the majority of up- or down-regulated genes but a sub-set of up- and down- regulated genes show a reduction in copy number (Fig. 5C).

Taken together, the mutants displayed unique gene expression profiles with mutant 9.7 showing differential gene expression of genes associated with amino acid catabolism while mutant 11.1 upregulates several genes associated with amino acid, sugar transported as well as for heat shock proteins.

## Discussion

*S. pastorianus* comprises a set of strains with interesting fermentative traits, including the ability to utilise complex sugars such as maltotriose, to produce aromatic compounds above the sensory threshold, and to ferment at cold temperatures (1, 3, 25-28). Their complex aneuploid genomes underpin these unique physiological traits. The hybrids, formed just 500-600 years ago, appear to still be in genomic flux as evidenced from copy number variations in published sequences of strains (4, 5, 12, 29). Additionally, CNVs have been observed during a single round of fermentation conducted in high gravity wort at 20°C (23). Indeed, we observed homologous recombination between two known recombination sites on chromosome VII in two out of thirty-six cultures in this study.

The sterile nature of the hybrids creates an evolutionary bottleneck for genetic variation. Approaches such as hybridisation, adaptive evolution, and allele replacement, have been used to generate strains with altered phenotypes such as temperature tolerance, sugar utilisation and aroma production (16). While successful, such approaches are time-consuming, hybrids can be unstable and require lengthy successive propagations to stabilise while adaptive evolution can require propagation for 300-400 generations to obtain successful phenotypes. Allele replacement is avoided due to legislation on the use of GMO products and customer rejection of GMO products (30).

Here we have established an accelerated evolution approach to obtain lager yeasts with altered flavour profiles through the temporary inhibition of the cells stress response by applying a severe short heat stress or by treatment of cells with radicicol, a Hsp90p, inhibitor. Both treatments have previously been shown to induce aneuploidy and/or chromosomal recombination (22, 23). Resistance to amino acid analogues of phenylalanine was used to select strains with an increased flux towards the production of higher alcohols and esters. Our results show that radicicol treatment, followed by selection on PFPA produced the greatest number of mutants. Mutants resistant to either amino acid analogue displayed cross resistance, but PFPA was more toxic to the cells than B2TA (31).

### Chromosomal Changes

Hsp90p assists in the folding of an extensive list of clientele proteins (32, 33). Amongst this list are proteins involved in DNA repair, kinetochore assembly and cell cycle checkpoint monitoring. The chromosomal changes identified in the mutants, aneuploidy, homeologous chromosome recombination and translocations can be accounted for by the temporary inhibition of chaperone function of Hsp90p coupled with the selection of clones resistant to amino acid analogues. There appears to be a correlation between the degree of resistance to PFPA and the extent of induced aneuploidy: mutant 9.7 from W34/70 is resistant to 400 µg · mL^-1^ PFPA and showed extensive aneuploidy while mutant 11.1 from CBS1538 was only resistant to 100 µg·mL^-1^ of PFPA and showed less induced aneuploidy. The difference in induced aneuploidy may also be a consequence to the tolerance of the strains to aneuploidy as CBS1538 is triploid with 48 chromosomes while W34/70 is almost pentaploid at 76 chromosomes.

There appears to be a preferential loss of *Se* over *Sc* chromosomes in the mutants. Previous analysis of chromosome composition following *de novo* generated hybrids of *S. cerevisiae* x *S. eubayanus* observed a similar preferential loss of *Se* chromosomes (18). The reason for this preferential loss is not currently understood but may reflect a greater fitness of strains with higher *Sc* content, or the essential requirement for some *Sc* alleles. We previously reported that *Sc* alleles are overrepresented in genes upregulated during fermentation in CBS1538 despite the extensive loss of *Sc* chromosomes in this strain (12). Interestingly, mutant 9.7 has reverted to a near tetraploid (n = 65) and there is some reciprocity in sub-genome loss to create the preferred stable ploidy for lager yeasts.

Previous studies showed that diploid *S. cerevisiae* strains treated with radicicol acquired an extra copy of chromosome XV, increasing the copy number of *STI1*, a Hsp90 co-chaperone and the multidrug resistance gene, *PDR5*, which are located on that chromosome. Here, we did not see increased copies of chromosome XV, most likely due to differences in selection pressures. Interestingly, we did observe a constitutive over expression of *Se HSP82*, which encodes for Hsp90p, in mutant 11.1 in all three conditions examined here. The heat shock genes *HSP104, HSP30, HSP26* and *SSA4*, encoded from the *Se* sub-genome, are also upregulated in mutant 11.1 on Day 2 in wort but these heat shock genes are downregulated in the mutant 9.7. Conversely, genes encoding Hsp78p and its co-chaperone Hsp42p, are upregulated in mutant 9.7 in wort on days 2 and 4. Taken together, it appears that the mutants display altered stress responses perhaps as a compensatory response to the imposed stress on the cells.

### SNPs in the mutants

Surprisingly, we detected few SNPs that were unique to the mutant strains, however crucially amongst those found were SNPs in *ARO4* in both mutants. Aro4p catalyses the first step of the aromatic amino acid biosynthetic pathway, the Shikimate pathway. The activity of Aro4p is inhibited by negative feedback of phenylalanine, tyrosine, and tryptophan. The mutations identified in mutants 11.1 and 9.7 are located in amino acid positions known to be involved in the negative feedback inhibition (34). Gene expression analysis in the mutants corroborates the *in situ* increase in aromatic amino acid biosynthesis as we observed the upregulation of *ARO9* and *ARO10*, two genes that are positively regulated by *ARO80* in the presence of increased concentrations of phenylalanine. Increased production of aromatic amino acids in the cell also drives the flux through the Ehrlich pathway towards the production of higher alcohols and esters such as 2-phenylethanol, 2-phenylethyl acetate, tryptophol, and tyrosol, observed in the mutants. Allele variation in *ARO4* may also contribute to the overproduction and the variation of aromatic compounds between the mutant strains. In mutant 9.7, it is the *Sc* copy of *ARO4* that is affected with a SNP present in two of the three copies of the gene while in mutant 11.1, the mutation occurs in the *Se* copy of *ARO4* with one of the three copies of the allele affected. The overproduction of 2-phenylethanol and 2-phenylethyl acetate by the yeast strains have a positive impact in the final beer as these two molecules imparts notes of honey and rose-like flavours. Furthermore, tyrosol and tryptophol contribute to in-mouth sensory properties of beverages (35-37).

The SNP analysis also uncovered allele frequency changes and loss of heterozygosity in the two variant *S. cerevisiae* sub-genomes, confirming the presence of two types of *Sc* chromosomes in the Group II strains (4). Additionally, we uncovered recombination between the *Sc1* and *Sc 2* chromosomes that has not previously been reported. Such recombination events may contribute to the genetic variation within *S. pastorianus* strains.

### Gene expression changes in the mutants

In addition to the notable upregulation of *ARO9* and *ARO10* in mutant 9.7 and heat shock genes in mutant 11.1, several additional transcriptome changes were observed in the mutant strains. Of note was the upregulation of *BAP2* and *BAT2*, encoding for a branched chain amino acid permease and branched chain amino acid transaminase and the ammonium permease *Se MEP2* in mutant 11.1. Both *Sc* and *Se MEP1* are upregulated in mutant 9.7. Such upregulation increases the uptake of nitrogen and amino acids into the cell. Genes involved in lysine and methionine metabolism are also upregulated in mutant 9.7. Taken together these changes improve the fluxes toward the production of higher alcohols and esters in the mutants.

The significant loss of chromosome copies in mutant 9.7 accounts for most downregulated genes. Inversely, genes located in chromosomes with a higher copy number show significant higher expression. This confirms our previous finding that gene expression is directly correlated with gene copy number in *S. pastorianus*. Such alterations in the ratios of *Sc* and *Se* alleles may increase proteome diversity through generating chimeric multi sub-unit complexes with altered composition of sub-units thus affecting interactions with substrates.

In summary, the present work confirms that the plasticity of *S. pastorianus* genomes is an advantageous trait of these strains supporting opportunities for genetic diversity in otherwise sterile strains.

## Methods

### Yeast strains and growth conditions

CBS1538, was obtained from the Collection de levures d’intérêt biotechnologique, Paris, France and Weihenstephan 34/70 (W34/70) was kindly supplied by Dr. Jurgen Wendland, Geisenheim Hoch Universitat, Germany. Strains were grown in YPDM (1% (w/v) yeast extract, 2% (w/v) peptone and 1–2% (w/v) of both dextrose and maltose at 20–25°C or in minimal medium (0.17% Yeast Nitrogen Base (YNB) w/o amino acids and ammonium salts supplemented with 1% of dextrose and 1% of maltose, and 0.5% (NH_4_)_2_SO_4_ as a nitrogen source at 20°C.

Small-scale fermentations (10mL) and large-scale fermentations (2L) were carried out in 12% wort containing 1mM ZnSO_4_ in 15 mL glass test tubes or 3L tall tubes in triplicate or duplicate, respectively. Cells were propagated in 4% YPDM at 25ºC and pitched at a cell density of 1.5×10^7^·mL^-1^. The tubes were fitted with a water trap airlock attached to a bung and incubated at 13ºC at a 45° angle without shaking. The specific gravity of the wort was measured using a refractometer. The residual cultures from the small-scale fermentations were centrifuged and the cells saved for RNA extraction.

For the radicicol treatment, Synthetic Complete medium (SC;1% of dextrose, 1% of maltose, 0.5% of (NH_4_)_2_SO_4_, 0.17% of YNB w/o amino acids and ammonium salts and 0.2% of a mix of amino acids was used.

### Mutagenesis

For the Heat Shock Thermal Stress (HSTS), 5 mL of cultures at 1×10^7^ cells mL^-1^ were heated at 45-55°C for 10-15 minutes. After a recovery of 3-5 h, aliquots of 100 µL were plated onto minimal media agar plates containing amino acid analogues β-(2-thienyl)-DL-alanine (B2TA) or p-fluorophenylalanine (PFPA) at concentrations 75 µg mL^-1^ or 200 µg mL^-1^ respectively.

For radicicol treatment, a modification of the method proposed by (22) was followed. Cells at a concentration of 1×10^7^ cells·mL^-1^ were inoculated in SC medium containing a final concentration of radicicol of 20, 40 and 100 µg · mL^-1^ for 24 and 48h. After the treatment, aliquots of 100 µL were plated onto minimal medium agar plates containing 75 µg·mL^-1^ of B2TA or 200 µg· mL^-1^ of PFPA. To ensure that mutants retained the phenotype of resistance to amino acid analogues, the colonies were re-streaked onto agar plates of minimum media containing the amino acid analogues at least three times. Resistant colonies were designated by a code based on the mutagenesis treatment and conditions (*SI Appendix*, Table S1).

### GC-MS analysis

Analysis of volatile compounds in wort at the end of fermentation were conducted as previously described (38) with the following modifications. Briefly, 0.5 mL of the samples in 20 mL vials were supplemented with sodium chloride to a final concentration of 1g · mL^-1^ and 25 μL of the 2-octanol as the internal standard (final concentration 200 μg · L^-1^). All samples were incubated for 10 min at 40°C, then the volatile compounds were collected on a divinylbenzene/carboxen/polydimethylsiloxane fibre (DVB-CAR-PDMS) coating 50/30, and 2-cm length SPME fibre purchased from Supelco (Sigma Aldrich, Milan, Italy) for 40 min. GC analysis was performed on a Trace GC Ultra gas chromatograph coupled with a TSQ Quantum Tandem mass spectrometer (Thermo Electron Corporation, USA) (47). Identification of compounds was based on comparison with a mass spectral database (NIST version 2.0) and with the retention time of the reference standards. For the screening experiment, semi-quantitative results of the interested volatile organic compounds (VOCs) were calculated. The relative amount of each volatile was expressed as µg · L^-1^ of 2-octanol (48). For the scaling-up confirmation experiment, interested VOCs were quantified. Calibration curves were measured with standards dissolved in simulated beer solution (7% ethanol in water, v/v).

### DNA extraction, *de novo* genome sequencing

DNA extraction was carried out as described by Querol, Barrio and Ramón (39). *De novo* genome sequencing was carried out by Novogene (www.novogene.com) with Illumina technology on paired-end reads (150bp). The genome sequences are available for deposition into the SRA database at the National Centre for Biotechnology Information. After trimming paired-end reads, the reads were mapped with bowtie2 (version 2.4.2) (40) to the parental genomes of *S. pastorianus* CBS1483 or to a combined genome of *S. cerevisiae* and *S. eubayanus* derived from *S. cerevisiae* S288C, assembly R64 and *S. eubayanus* (SEU3.0). The annotation of the *S. eubayanus* genome was updated by blasting against the *S. cerevisiae* annotation in assembly R64. High quality matches sharing >50% identity and >75% of the coverage of the *S. eubayanus* protein length was accepted as orthologs and renamed as alleles in the annotation. Of the 805 orthologs identified, 672 had >70% identity.

### qPCR

To confirm copy number variations, quantitative PCR using specific primers to discriminate between sub-genomes (SI Appendix, Table S8) was conducted as previously described (41).

### Copy number variation and SNP analysis

Reads from *de novo* DNA sequencing, mapped against the combined genomes of the reference strains of *S. cerevisiae* and *S. eubayanus* were transformed into sorted BAM files using samtools and the data was extracted as reads/500 bp and normalized by the size of the library (total number of reads) to determine chromosome copy number.

Samtools were used to call SNP/InDel from BAM files. ANNOVAR was used to annotate variants (42). A gene-based annotation file was generated using the genome of CBS1483 and a file containing the transcriptome of the same strain using the ANNOVAR tools. Synonymous, non-synonymous, stop gain and stop loss SNPs common to both the wildtype and mutant strains were called. Non-synonymous SNPs located in exonic regions and unique to the mutant strains were selected and any with a quality score of <100 and a depth read coverage score of <50, and with an incorrect allele frequency according to the gene copy number were discarded. The remainder were manually verified using Integrated Genome Viewer (IGV_2.11.0) (43).

### RNA extraction, sequencing, and analysis

RNA extraction was carried out as described by de la Cerda, *et al*. (12). RNA sequencing was conducted on cDNA libraries using the HiSeq4000 Illumina Platform at the Genomic Technologies Core Facility at the University of Manchester. The libraries were generated using the TruSeq Stranded mRNA assay (Illumina, Inc.) according to the manufacturer’s protocol. The RNAseq dataset is available for deposition to the GEO database at NCBI

Read counts from the RNA mapping were uploaded into iDEP9.1 (44) and analysed as previously described de la Cerda (2022). RNA from 2 cultures of the mutant strain 9.7 were omitted from further analysis due to the observation of a recombination event on chromosome VIII between *XRN1* and *ZUOI*. Differentially expressed genes were manually removed if they had zero coverage in either strain or had read counts <20. Gene ontology enrichment was carried out using ClueGO (45) with the following parameters, p-value <0.05, minimum of 3 genes per pathway, and a Kappa score of 0.4. A two-sided hypergeometric test and Bonferroni step-down pV correction were applied.

## Acknowledgements

We thank the Genomic Technologies Core Facility and the Bioinformatic Core Facility at the University of Manchester for assistance with RNAseq analysis and Ian Donaldson for initial bioinformatic support. We thank all the members of Project Aromagenesis @aromagenesis for their stimulating scientific discussions and their support during the project. RC was supported by European Union’s Horizon 2020, Marie Skłodowska Curie Innovative Training Networks Programme grant number 76436.

## Supplementary Figures

S1. Volatile profiles of CBS1538 and W34/70. Aromatic profiles of the parental strains CBS1538 and W34/70. Black columns,CBS1538, hatched columns,W34/70. Error bars represent the standard deviations from the mean of duplicate fermentations. Compounds showing statistical differences between the mutants and the parental strains are indicated by connecting lines, p≤0.001,***; p≤0.01,**, p≤0.05,*.

S2. Confirmation of aneuploidy of 9.7 compared to W34/70 by qPCR. Error bars represent the standard deviations from the mean of three independent experiments.

S3. Chromosome changes observed between two Sc sub-genomes. A) SNP observed in *ScARO4*, where mutant obtained a new allelic variant in one chromosome, followed by a chromosome loss of the wildtype allele and a chromosome gain of the mutant allele. B) Loss of Heterozygosity observed in *ScSTI1, ScCIN5* and intergenic regions in mutant 9.7.

S1 Table. Number of mutants obtained after several rounds of evolution using HSTS and radicicol. S2 Table. Non-synonymous SNPs found in mutants 11.1 and 9.7.

S3 Table. Raw read counts: *S. pastorianus* genome mapped against combined *S. cerevisiae* and *S. eubayanus* genomes.

S4 Table. Differentially expressed genes observed in mutants compared to their respective parental strains.

S5 Table. Venn diagram lists of up- or downregulated genes in mutant 9.7. S6 Table. Venn diagram lists of up- or downregulated genes in mutant 11.1.

S7 Table. Lists of genes associated with differentially express gene ontologies in mutant 9.7.

S8 Table. Primers used in this study.

