## Supplementary figures and images for "Enhanced flavour profiles through radicicol induced genomic variation in *S. pastorianus* lager yeast"

### SI Appendix, Figures

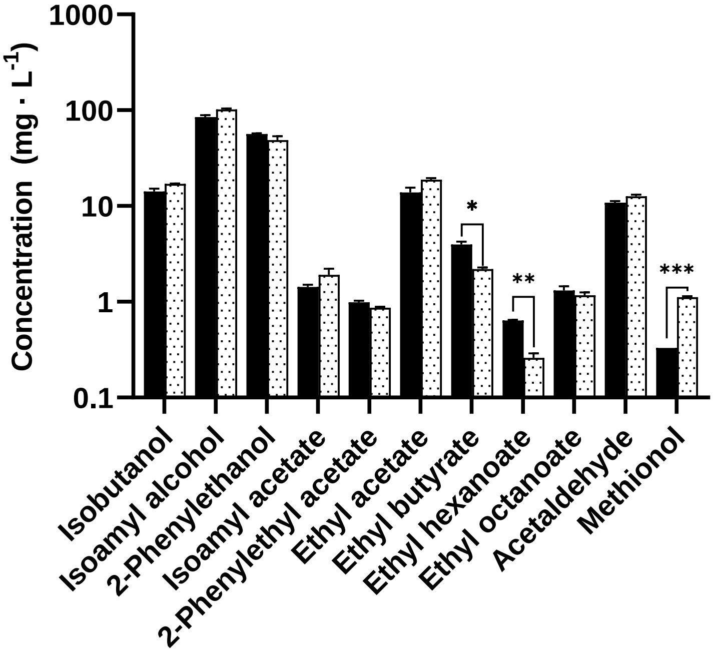


**SI Appendix, Fig S1**

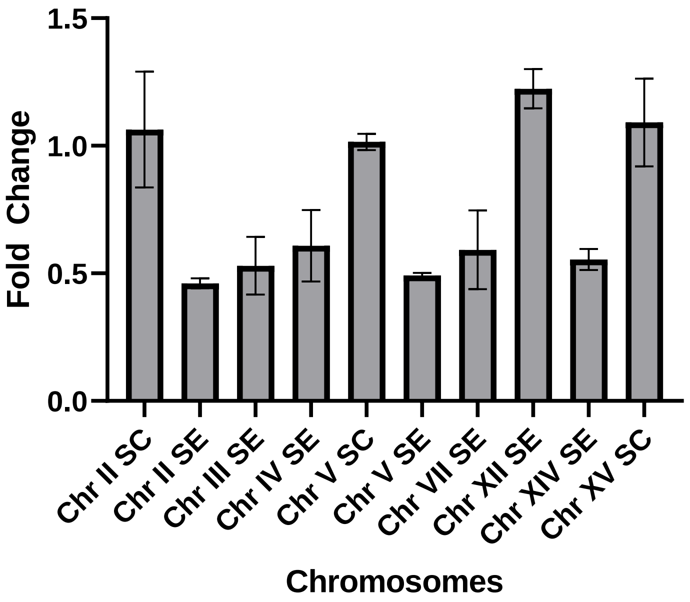


**SI Appendix, Fig S2**

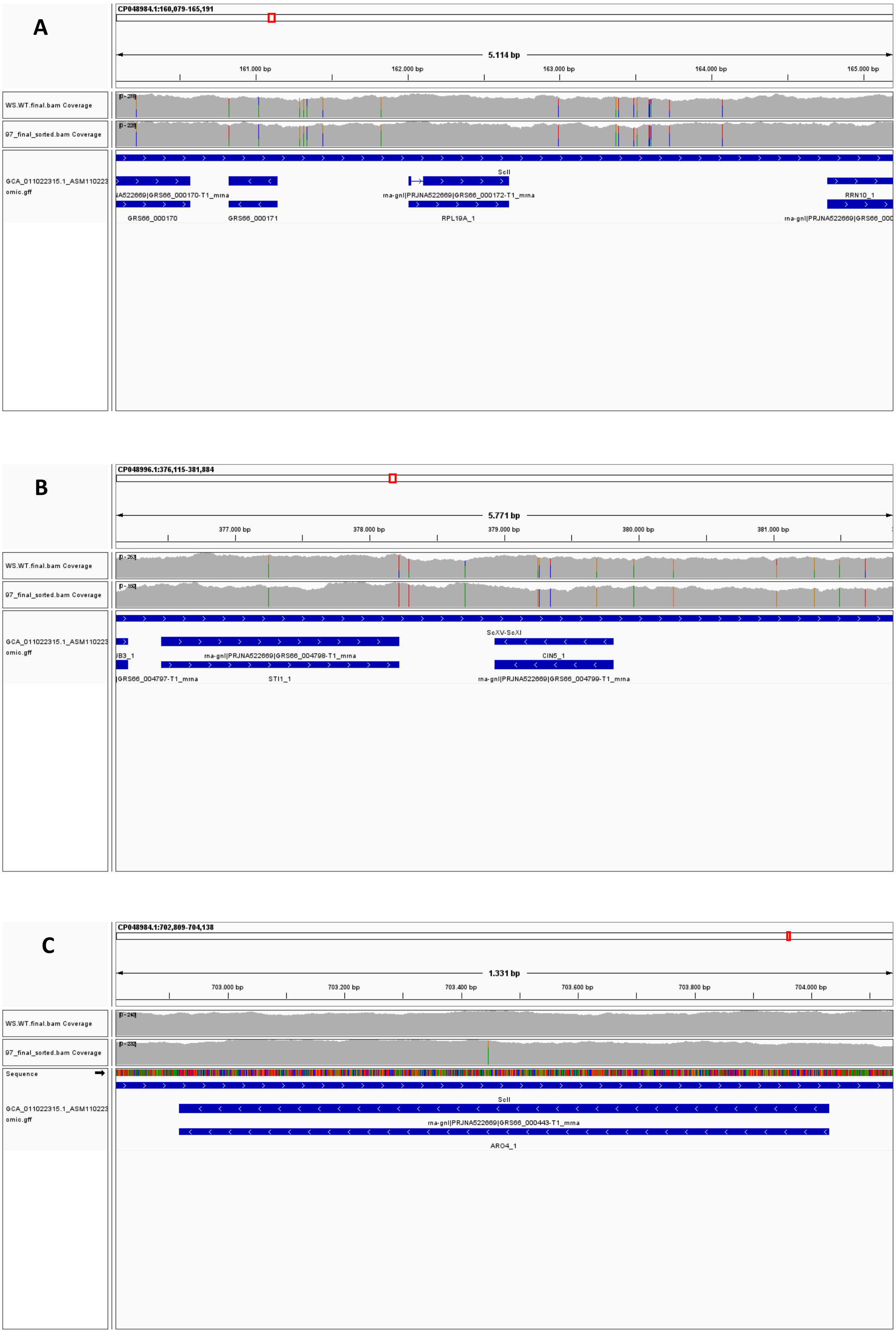


SI Appendix, Fig S3
